# Adhesion-clutch drives three-dimensional axon outgrowth

**DOI:** 10.64898/2026.09.07.749764

**Authors:** Hongjia Chen, Zhen Qiu, Saki Yamada, Akinori Murakami, Takunori Minegishi, Naoyuki Inagaki

## Abstract

Axon outgrowth requires forces generated by the growth cone. A key model explaining this force generation is the adhesion-clutch mechanism, in which adhesion and clutch molecules convert backward movement of actin filaments into the force that drives axon outgrowth. However, this mechanism has not been validated in three-dimensional (3D) environments. Additionally, a recent study reported that inhibiting actin dynamics or the cell adhesion molecule integrin did not affect axon outgrowth in a 3D collagen gel, challenging the adhesion-clutch paradigm. Here, we show that the adhesion molecule N-cadherin and the clutch molecule shootin1a form a non-integrin adhesion-clutch in a 3D environment containing an appropriate adhesive substrate, N-cadherin. We detected forces produced by growth cones when N-cadherin was present. Furthermore, inhibition of N-cadherin, shootin1a or actin dynamics suppressed 3D axon outgrowth. Our findings demonstrate that the adhesion-clutch is critical machinery for 3D neural network formation under the regulation of specific adhesions.

## INTRODUCTION

Axon outgrowth is an essential step of neural network formation. The growth cone, discovered by Santiago Ramón y Cajal in 1890^1^, is a cytoskeletal-rich structure located at the tip of extending axons. It senses extracellular chemical and mechanical cues to guide axon outgrowth^2–6^. Remarkably, growth cones, even when separated from the neurite shaft, continue to move forward in a two-dimensional (2D) environment^7^, suggesting their intrinsic ability to generate driving forces for axon outgrowth. A subsequent study in 2D environments confirmed this by demonstrating that growth cones pull the axonal shaft via adhesion^8^. Analyses using traction force microscopy further showed that growth cones generate traction forces at the levels of 1-40 Pa (= pN/µm^2^)^9–11^.

A key model explaining this force generation is the adhesion-clutch mechanism^12,13^. Actin filaments (F-actins) undergo continuous retrograde flow in growth cone^14–16^, driven by their polymerization at the leading edge and proximal disassembly together with myosin II activity^16^ (Figure S1A, left). Cell adhesion and clutch molecules transmit the movement of the F-actin flow to the adhesive extracellular environment, converting it into the force that drives axon outgrowth^17–19^ (Figure S1A, right). However, this mechanical transmission has not yet been demonstrated in a 3D environment. In fact, a recent study reported that F-actin disruption or inhibition of the adhesion molecule integrin had no effect on axon outgrowth in 3D collagen gels^20^. Additionally, the force produced by growth cones was undetectable in this environment. These data challenge the adhesion-clutch paradigm, particularly in physiological 3D environments^20,21^.

N-cadherin is a cell adhesion molecule that plays critical roles in neuronal development^22^. N-cadherin localized at the neuronal plasma membrane interacts homophilically with N-cadherin expressed on neighboring cells^22,23^. In 2D environments, N-cadherin coated on the adhesive substrate promotes axon outgrowth^18,24^, where N-cadherin in the growth cone membrane links F-actin retrograde flow to the substrate for force generation^18^. On the other hand, shootin1a (*SHTN1*) is a neuronal clutch molecule^17^. It mediates an adhesion-clutch for axon outgrowth and guidance in 2D conditions^17,25^ by linking the F-actin flow to the cell adhesion molecules L1^17^ and receptor deleted in colorectal cancer (DCC)^26^.

Here, we demonstrate that N-cadherin and shootin1a form an adhesion-clutch when their specific adhesive ligand, N-cadherin, is present in 3D environments. We also detected the traction force produced by growth cones. Furthermore, inhibition of N-cadherin/shootin1a or F-actin disruption suppressed the 3D axon outgrowth in the presence of extracellular N-cadherin. Our findings demonstrate that the adhesion-clutch mechanism is fully operational for 3D axon outgrowth when suitable adhesion is engaged.

## RESULTS

### N-cadherin and shootin1a form adhesion-clutch for 2D axon outgrowth

N-cadherin on neighboring cells acts as an adhesive ligand of N-cadherin localized at the neuronal plasma membrane^22,23^. To examine their role in axon outgrowth, we cultured mouse hippocampal neurons on glass coverslips coated sequentially with polylysine and N-cad-Fc (chimeric protein containing the extracellular domain of N-cadherin and the Fc domain of IgG)^27^ or polylysine alone as a control substrate. N-cad-Fc mimics N-cadherin on neighboring cells as the adhesive ligand. As reported previously^18,24^, N-cad-Fc coated on the substrate promoted 2D axon outgrowth (Figure 1A). Additionally, knockdown of neuronal N-cadherin (Figure S1B) inhibited axon outgrowth on the N-cad-Fc-coated substrate (Figures 1B and S1C) (see also SUPPLEMENTAL DISCUSSION 1), indicating that the adhesion between the neuronal N-cadherin and its ligand N-cadherin mediates axon outgrowth.

**Figure 1.**
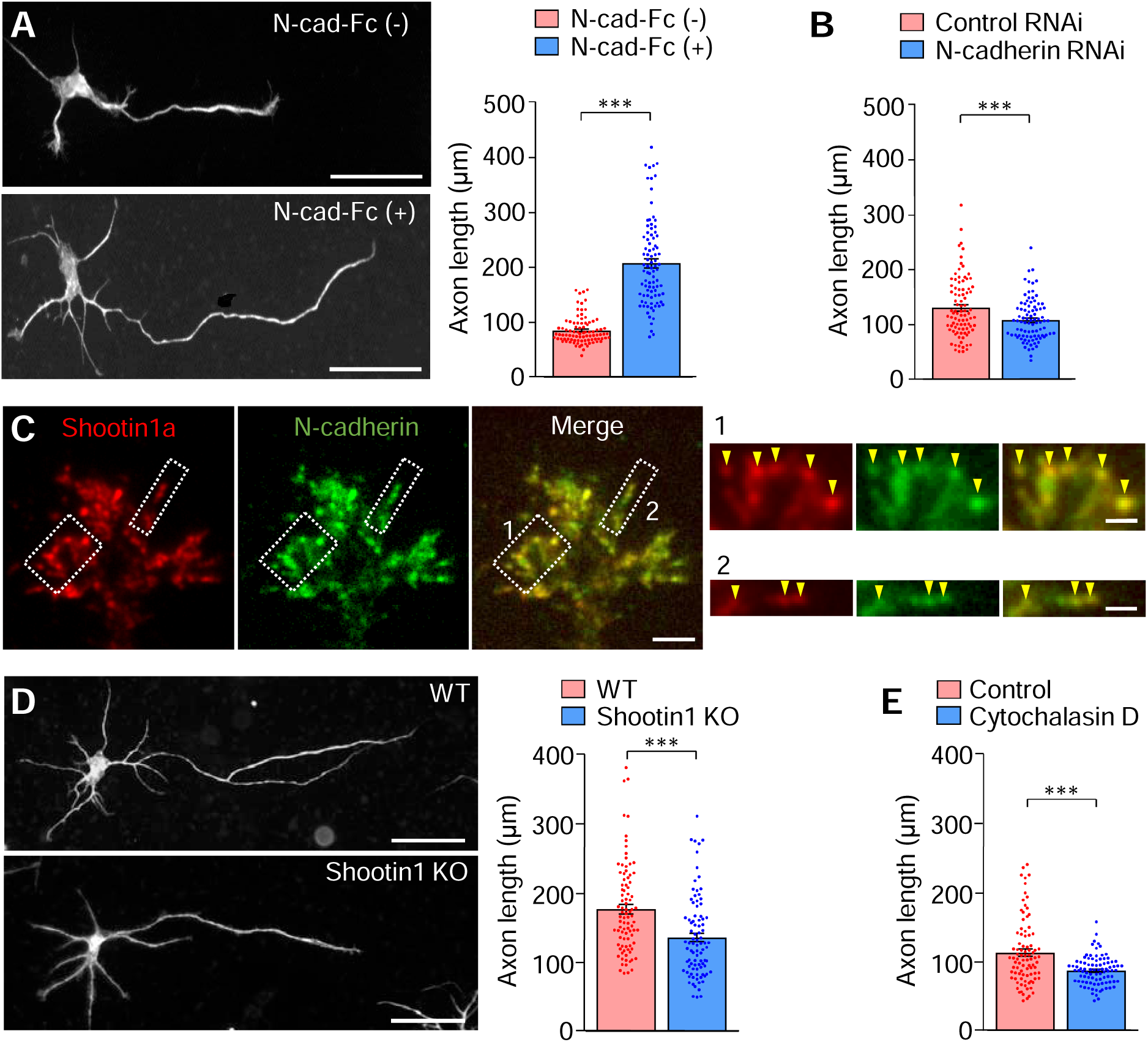
N-Cadherin and shootin1a promote 2D axon outgrowth. (A) Fluorescence images of days in vitro (DIV) 2 hippocampal neurons cultured on glass coverslips coated with polylysine alone (N-cad-Fc (-)) or coated sequentially with polylysine, anti-Fc and N-cad-Fc (N-cad-Fc (+)). Cells were immunostained with anti-Tuj1 antibody. Quantification of axon length is shown to the right. Mann-Whitney *U*-test; p < 1.0 × 10^−15^ (N-cad-Fc (-): n = 90 cells; N-cad-Fc (+): n = 90 cells). Scale bar, 50 μm. (B) Quantification of axon length of neurons transfected with control miRNA or N-cadherin miRNA (#1) and cultured on N-cad-Fc-coated coverslips shown in Figure S1C. Mann-Whitney *U-*test; p = 0.0052 (control miRNA: n = 90 cells; N-cadherin miRNA (#1): n = 90 cells). Scale bar, 50 μm. (C) Fluorescence images of axonal growth cone of DIV 2 hippocampal neurons double-immunostained with anti-shootin1a (red) and anti-N-cadherin-ICD (green) antibodies. Enlarged views of the lamellipodium in white square 1 and the filopodium in white square 2 are shown to the right. Arrowheads indicate the co-localization between shootin1a and N-cadherin. Scale bar, 5 μm (in the enlarged images, 2 μm). (D) Fluorescence images of DIV 2 hippocampal neurons derived from WT or shootin1 KO mice and cultured on N-cad-Fc-coated coverslips. Cells were immunostained with anti-Tuj1 antibody. Quantification of axon length is shown to the right. Mann-Whitney *U*-test; p = 3.1 × 10^−6^ (WT: n = 90 cells; shootin1 KO: n = 90 cells). Scale bar, 50 μm. (E) Quantification of axon length of DIV 2 hippocampal neurons seeded on N-cad-Fc-coated coverslips and treated with 0.1% dimethyl sulfoxide (DMSO, control) or 1.0 μM cytochalasin D shown Figure S1E. Mann-Whitney *U*-test; p = 1.3 × 10^−4^ (DMSO: n = 90 cells; Cytochalasin D: n = 90 cells). Data represent means ± SEM. ∗∗∗p < 0.01.

We previously reported that the clutch molecule shootin1a directly interacts with the intercellular domain (ICD) of N-cadherin^28^. Shootin1a colocalized with N-cadherin in axonal growth cones (Figures 1C and S1D). Furthermore, shootin1a knockout (KO) or F-actin disruption by 1 μM cytochalasin D inhibited axon outgrowth on N-cad-Fc-coated substrate (Figures 1D, 1E and S1E) (see also SUPPLEMENTAL DISCUSSION 2), indicating that shootin1a and F-actins are involved in N-cadherin-mediated axon outgrowth.

The engagement of adhesion-clutch (i.e., the coupling between F-actin retrograde flow and the adhesive substrate) mechanically impedes the F-actin flow, and generates traction force by transmitting the F-actin movement to the substrate (Figure S1A)^11,13,19^. To examine whether N-cadherin and shootin1a constitute adhesion-clutch for axon outgrowth, we next monitored F-actin retrograde flow within growth cones and traction force produced by growth cones. F-actin retrograde flow was visualized by fluorescent speckle imaging of HaloTag-actin, where the fluorescently labeled actin molecules replace some of the endogenous actins to form F-actins^29^. HaloTag-actin speckles moved retrogradely in growth cones (Video S1), depicting the F-actin movement (Figure 2A). The speed of F-actin flow on N-cad-Fc was 2.0 ± 0.1 μm/min and increased by removing N-cad-Fc (Figure 2A; Video S1) or by shootin1a (KO) (Figure 2B; Video S2). For force measurement, neurons were cultured on N-cad-Fc-coated or control polyacrylamide gels embedded with 200-nm fluorescent beads. Traction force generated by growth cones was analyzed by visualizing force-induced deformation of the gel, which is reflected by displacement of the beads from their original positions^29^. The beads moved toward the center of the growth cone (Figure 2C; Video S3). The magnitude of the traction force generated on N-cad-Fc was 17.5 ± 0.9 pN/μm^2^, and decreased by removing N-cad-Fc (Figure 2C; Video S3) or by shootin1a KO (Figure 2D; Video S4). Together, these data indicate that N-cadherin and shootin1a form an adhesion-clutch to drive 2D axon outgrowth.

**Figure 2.**
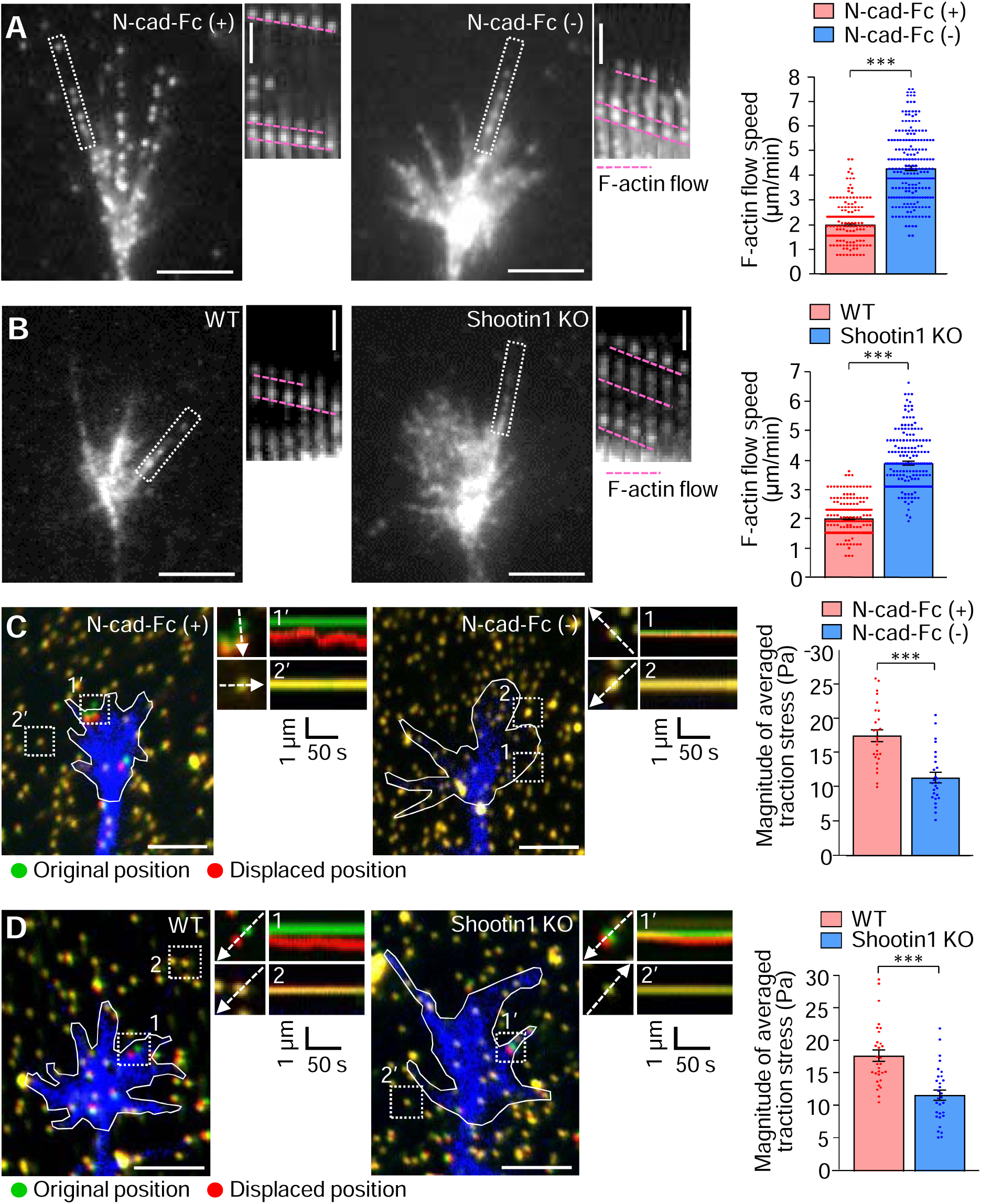
N-Cadherin and shootin1a form 2D adhesion-clutch. (A and B) Fluorescent speckle images of HaloTag-actin in axonal growth cones of DIV 2 hippocampal neurons. WT neurons were cultured on polylysine (N-cad-Fc (-)) or N-cad-Fc-coated (N-cad-Fc (+)) glass-bottom dishes (A). See Video S1. WT or shootin1 KO neurons were cultured on N-cad-Fc-coated glass-bottom dishes (B). See Video S2. Time-lapse montages of HaloTag-actin speckles in filopodia (boxed areas) obtained at 5-s intervals for 295 s are shown to the right. Pink dashed lines indicate the retrograde flow of speckles. Statistical analyses of F-actin flow speed are shown in the graphs on the right. Mann-Whitney *U-*test; p < 1.0 × 10^−15^ (A; N-cad-Fc (+): n = 207 speckles; N-cad-Fc (-): n = 238 speckles), p < 1.0 × 10^−15^ (B; WT: n = 210 speckles; shootin1 KO: n = 200 speckles). Scale bars, 5 μm (in the kymographs, 2 μm). (C and D) Fluorescence images showing axonal growth cones of DIV2 hippocampal neurons expressing EGFP (blue) and cultured on polyacrylamide gel embedded with 200-nm fluorescent beads. WT neurons were cultured on polylysine (N-cad-Fc (-)) or N-cad-Fc-coated (N-cad-Fc (+)) polyacrylamide gel (C). See Video S3. WT or shootin1 KO neurons were cultured on N-cad-Fc-coated polyacrylamide gel (D). See Video S4. The original and displaced positions of the beads in the gel are indicated by green and red colors, respectively. White lines indicate the growth cone boundaries. The kymographs along the axis of bead displacement (white dashed arrows) at the indicated areas show the movement of beads recorded every 3 s for 147 s. The beads in areas 2 and 2’ are reference beads. Statistical analyses of the magnitude of traction forces under axonal growth cones are shown in the graphs on the right. Two-tailed unpaired Student’s *t*-test; p = 3.0 × 10^−6^ (C; N-cad-Fc (-): n = 26 growth cones; N-cad-Fc (+): n = 26 growth cones), p = 3.1 × 10^−6^ (D; WT: n = 30 growth cones; shootin1 KO: n = 30 growth cones). Scale bars, 5 μm. Data represent means ± SEM. ∗∗∗p < 0.01.

### N-cadherin and shootin1a form 3D adhesion-clutch and generate traction force

Next, to examine whether N-cadherin and shootin1a mediate adhesion-clutch in a 3D environment, we cultured hippocampal neurons in 3D Matrigel^30^ in the presence or absence of the adhesive ligand N-cad-Fc. By fluorescent speckle imaging of HaloTag-actin, we detected retrograde flow of F-actins in the growth cones in this condition (Figure 3A; Video S5). The speed of F-actin flow in the presence of N-cad-Fc was 2.3 ± 0.1 μm/min and increased by removing N-cad-Fc or by shootin1a KO (Figure 3A; Video S5), thereby indicating that N-cadherin and shootin1a form the adhesion-clutch also in this 3D environment.

**Figure 3.**
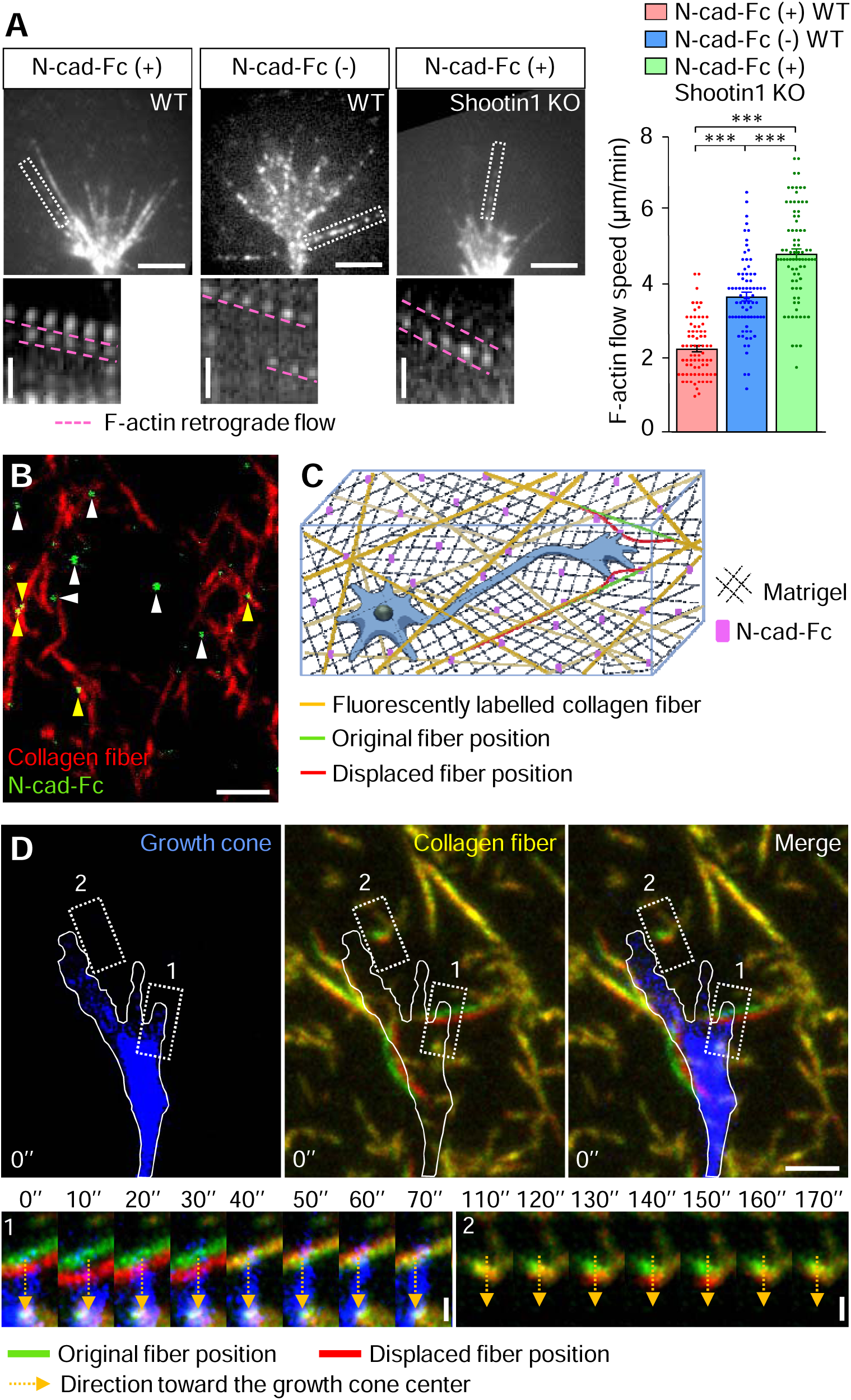
N-cadherin and shootin1a form 3D adhesion-clutch and generate traction force. (A) Fluorescent speckle images of HaloTag-actin in axonal growth cones of WT neurons (DIV 2) cultured in 3D Matrigel with (N-cad-Fc (+), WT) or without (N-cad-Fc (-), WT) the addition of N-cad-Fc, and shootin1 KO neurons (DIV 2) cultured in Matrigel containing N-cad-Fc (N-cad-Fc (+), Shootin1 KO). See Video S5. Time-lapse montages of indicated rectangular regions at 5-s intervals for 295 s are shown below. Pink dashed lines indicate F-actin retrograde flow. Statistical analyses of F-actin flow speed are shown in the graphs on the right. Two-tailed unpaired Welch’s *t-*test; p < 1.0 × 10^−15^ (N-cad-Fc (+) WT versus N-cad-Fc (-) WT), p < 1.0 × 10^−15^ (N-cad-Fc (+) WT versus N-cad-Fc (+) Shootin1 KO). Two-tailed unpaired Student’s *t*-test; p < 3.7 × 10^−9^ (N-cad-Fc (-) WT versus N-cad-Fc (+) Shootin1 KO). (N-cad-Fc (+) WT: n = 75 speckles; N-cad-Fc (-) WT: n = 76 speckles; N-cad-Fc (+) Shootin1 KO: n = 80 speckles). Scale bars, 5 μm (in the time-lapse montages, 2 μm). (B) Fluorescence image of Matrigel containing fluorescently labelled collagen fibers (red) and N-cad-Fc immunostained with anti-N-cadherin-ECD antibody (green). N-cad-Fc are localized in close proximity to collagen fibers (yellow arrowheads) and within Matrigel (white arrowheads). Scale bar, 2 μm. (C) A schema of traction force detection in 3D Matrigel containing fluorescently labelled collagen fibers (yellow lines) and N-cad-Fc (purple). The traction force generated by the axonal growth cone can be detected by monitoring force-induced fiber displacement (red line) from the original position (green line). (D) Fluorescence images showing an axonal growth cone of a DIV2 hippocampal neuron cultured in 3D Matrigel containing fluorescently labeled collagen fibers and N-cad-Fc. The growth cone was visualized by 5-chloromethylfluorescein diacetate (CMFDA) staining (blue). The original and displaced positions of collagen fibers are indicated by green and red colors, respectively. See Video S6. The white line indicates the boundary of the axonal growth cone. Dashed rectangle 1 shows a fiber directly attached to a filopodia; dashed rectangle 2 shows a fiber that is not directly attached to a filopodia. The time-lapse montages in the bottom panel show fiber movement in the indicated rectangular regions 1 and 2 (yellow dashed arrows), which were recorded every 10 s for 190 s. Scale bars, 2 μm (in the kymographs, 0.5 μm). Data represent means ± SEM. ∗∗∗p < 0.01.

Regarding the detection of forces generated in a 3D environment, we had difficulty preparing Matrigel that contain fluorescent beads because Matrigel mixed with fluorescent beads did not polymerize. Therefore, we utilized fluorescently labeled collagen fibers^31^. We prepared 3D Matrigel containing fluorescently labeled collagen fibers^32,33^ and N-cad-Fc (Figures 3B and 3C). Confocal laser microscopy demonstrated the localization of N-cad-Fc in close proximity to collagen fibers (yellow arrowheads, Figure 3B) and within Matrigel (white arrowheads). Due to the heterogeneous shape and elasticity of the fibers, it is not feasible to quantify the force as precisely as with a polyacrylamide gel. However, force generation can be detected through force-induced displacement of the fibers (red lines, Figure 3C)^31^.

Video S6 shows live imaging of a growth cone and surrounding collagen fibers. In 2D conditions, the traction force produced by the growth cone is very weak at 1-40 Pa (= pN/µm^2^)^9–11^ (Figures 2C and 2D), up to ∼100 times weaker than that produced by fibroblasts^9^. To detect weak forces, we focused on the movement of fibers perpendicularly oriented to the growth cone filopodia. We successfully detected the movement of collagen fibers attached to the growth cone filopodia toward the center of the growth cone (area 1, Figure 3D; Video S6). We also observed the centripetal movement of the fibers that were not directly attached to the filopodia (area 2, Figure 3D; Video S6), suggesting that Matrigel deformation induced by nearby filopodia causes the fiber movement indirectly. Together, we conclude that N-cadherin and shootin1a link F-actin flow in growth cones and the 3D environment, thereby generating traction force.

### N-cadherin-shootin1a adhesion-clutch drives 3D axon outgrowth

Finally, axon outgrowth in 3D Matrigel was analyzed. As observed in the 2D condition, hippocampal neurons actively extended axons in the presence of the adhesive ligand N-cad-Fc (Figure 4A). Importantly, inhibition of N-cadherin-shootin1a adhesion-clutch by removing N-cad-Fc (Figure 4A), shootin1 KO (Figure 4A), or N-cadherin knockdown (Figures 4B and S1F) reduced 3D axon outgrowth (see also SUPPLEMENTAL DISCUSSION 3). Furthermore, disruption of F-actin by 1 μM cytochalasin D inhibited axon outgrowth in 3D Matrigel in the presence of N-cad-Fc (Figure 4C). A previous study reported that disruption of F-actin by cytochalasin D had no effect on axon outgrowth in 3D collagen gel^20^. So, we also prepared 3D collagen gel in the presence of N-cad-Fc; confocal laser microscopy demonstrated N-cad-Fc attachment on the collagen fibers (Figure 4D). In this condition, F-actin disruption by 1 μM cytochalasin D inhibited axon outgrowth in 3D collagen gel (Figure 4E). However, as reported^20^, cytochalasin D had no effect on the axon outgrowth in the absence of N-cad-Fc (Figure 4F). Together, these data indicate that N-cadherin-shootin1a adhesion-clutch drives 3D axon outgrowth in response to its specific adhesive ligand.

**Figure 4.**
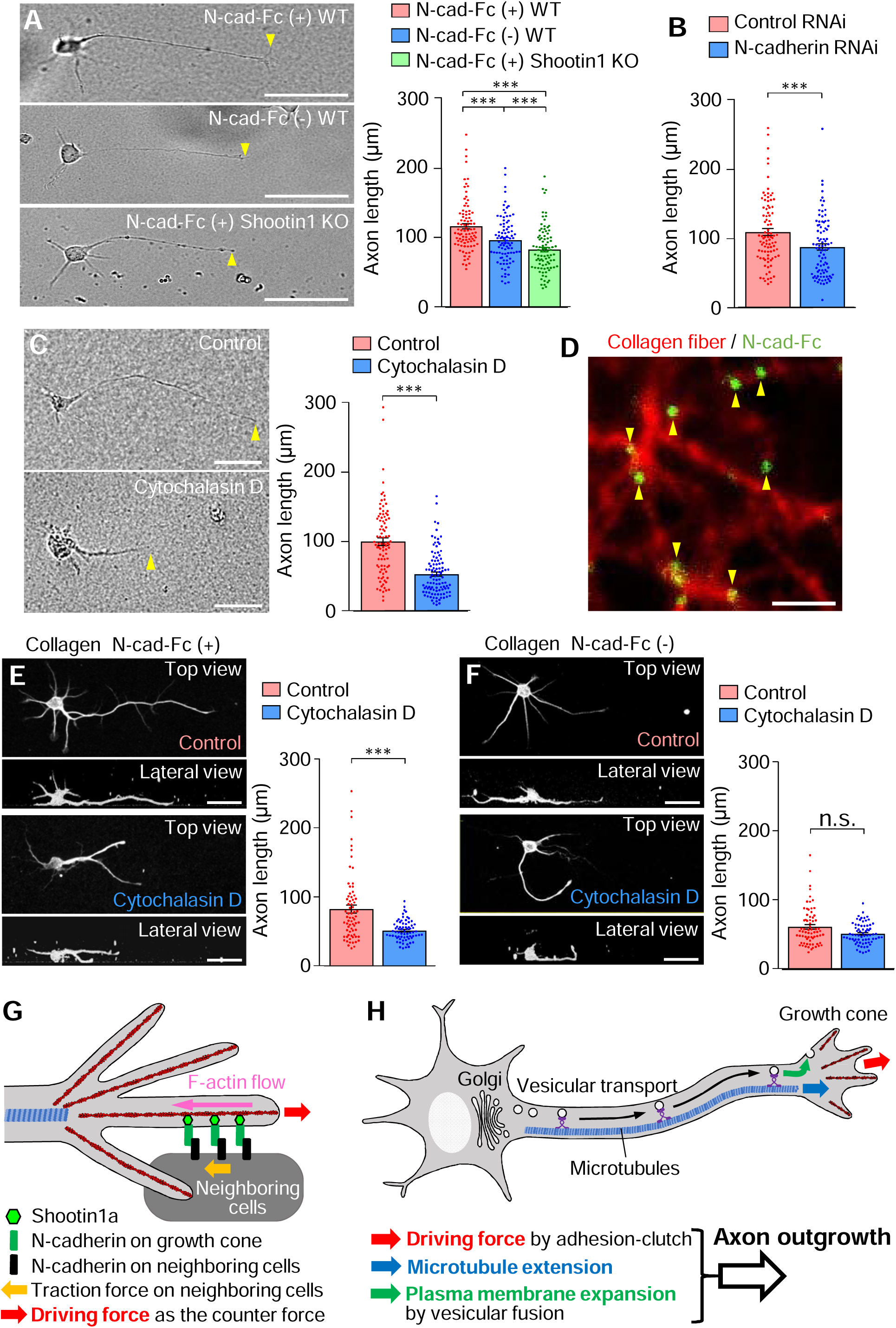
N-cadherin-shootin1a adhesion-clutch drives 3D axon outgrowth. (A) Brightfield images of DIV 2 hippocampal neurons cultured in 3D Matrigel with (N-cad-Fc (+), WT) or without (N-cad-Fc (-), WT) the addition of N-cad-Fc, and shootin1 KO neurons (DIV 2) cultured in Matrigel containing N-cad-Fc (N-cad-Fc (+), Shootin1 KO). Yellow arrowheads indicate axonal tips. Quantification of axon length is shown to the right. Mann-Whitney *U*-test; p = 2.1 × 10^−4^ (N-cadherin (+) WT versus N-cadherin (-) WT), p = 0.0041 (N-cadherin (-) WT versus N-cadherin (+) Shootin1 KO), p = 1.2 × 10^−10^ (N-cadherin (+) WT versus N-cadherin (+) Shootin1 KO) (N-cadherin (+) WT: n = 90 cells; N-cadherin (-) WT: n = 90 cells; N-cadherin (+) shootin1 KO: n = 90 cells). Scale bar, 50 μm. (B) Quantification of axon length of neurons transfected with control miRNA or N-cadherin miRNA (#1) and cultured in 3D Matrigel containing N-cad-Fc shown in Figure S1C. Mann-Whitney *U*-test; p = 0.0022 (control miRNA: n = 90 cells; N-cadherin miRNA (#1): n = 90 cells). (C) Brightfield images of DIV 2 hippocampal neurons cultured in 3D Matrigel in the presence of N-cad-Fc and treated with 0.1% DMSO (control) or 1 μM cytochalasin D for 40 h. Yellow arrowheads indicate axonal tips. Quantification of axon length is shown to the right. Mann-Whitney *U*-test; p = 5.0 × 10^−13^ (DMSO: n = 90 cells; Cytochalasin D: n = 102 cells). Scale bar, 50 μm. (D) Fluorescence image of fluorescently labelled collagen gel (red) containing N-cad-Fc (green). Yellow arrowheads indicate the attachment of N-cad-Fc on collagen fibers. Scale bar, 2 μm. (E and F) Fluorescence images of DIV 2 hippocampal neurons cultured in 3D collagen gel with (E) or without (F) the addition of N-cad-Fc. Neurons were treated with 0.1% DMSO (control) or 1 μM cytochalasin D for 40 h, and immunostained with anti-Tuj1 antibody. Quantification of axon length is shown to the right. Mann-Whitney *U*-test; p = 5.7 × 10^−6^ (E; control: n = 70 cells; cytochalasin D: n =70 cells), p = 0.068 (F; control: n = 70 neurons; cytochalasin D: n =70 neurons). Scale bars, 50 μm. (H) A diagram showing how axon outgrowth is driven by 3D adhesion-clutch. At the leading edge of the axonal growth cone, the adhesion clutch, involving shootin1a, N-cadherin, and its adhesive ligand N-cadherin on neighboring cells, transmits the rearward movement of F-actin (pink arrow) to the neighboring cells. This produces a backward traction force (yellow arrow). The driving force of growth cone advance (red arrow) is produced as the counterforce of the traction force. (I) An integrated model for axon outgrowth. The adhesion-clutch provides the driving force that extends and guides axons (red arrow). Microtubule extension through polymerization (blue arrow) and plasma membrane expansion through vesicular insertion (green arrow) extend the axonal shaft. These processes cooperatively achieve axon outgrowth. Data represent means ± SEM; ∗∗∗p < 0.01, ns, not significant.

## DISCUSSION

### Adhesion-clutch drives 3D axon outgrowth

The present study demonstrates that N-cadherin and shootin1a form an adhesion-clutch in a 3D environment in the presence of the adhesive ligand N-cadherin. The traction force of the growth cone was also detected. Reduction of N-cadherin or shootin1a inhibited 3D axon outgrowth in the presence of N-cadherin. These findings, along with the previous reports that the inhibition of N-cadherin or shootin1 leads to defects in axon outgrowth and pathfinding *in vivo*^25,34–37^, indicate that the adhesion-clutch drives axon outgrowth in 3D environments (Figure 4G).

On the other hand, a previous study reported that F-actin disruption or integrin inhibition have no effect on axon outgrowth in 3D collagen gel^20^. Collagen is an adhesive ligand for integrins, not cadherins. Therefore, the functionality of an adhesion-clutch system depends on its constituent molecules and the appropriate ligands. Consistently, previous studies reported that the 3D ameboid migration of leukocytes depends on the adhesion-clutch system involving shootin1 and the adhesion molecule L1^38^, rather than integrin^39,40^.

### Cooperation of adhesion-clutch with microtubule and membrane dynamics

While the adhesion-clutch provides the necessary driving force, it also cooperates with other processes to achieve axon outgrowth. Axon outgrowth requires the extension of the axonal shaft, which is accompanied by an increase in the cytoplasmic volume and plasma membrane expansion^41,42^. Tubulin, the most abundant protein in the neurite shaft^43^, forms microtubules, and its polymerization and microtubule stabilization are essential for axon outgrowth^44,45^. In addition, plasmalemmal precursor vesicles are generated at the endoplasmic reticulum and Golgi apparatus in the cell body, transported along growing axons, and inserted at the growth cone membrane to expand axonal plasma membrane^41,42^. These processes are also required for axon outgrowth *in vitro* and *in vivo* (Figure 3H)^46–49^. Recent studies have shown that the application of weak pulling forces on axons stabilizes microtubules^50^ and enhances axon outgrowth^51^. Additionally, neurite extension increases the plasma membrane tension^52^, which in turn promotes membrane exocytosis^53^. Future studies are needed to determine whether the mechanical pull exerted by the adhesion-clutch machinery integrates these processes to promote axon outgrowth.

### Axonal pathfinding through adhesion between the growth cone and environment

Axon outgrowth and pathfinding *in vivo* are regulated by various extracellular cues, including soluble chemicals (chemotaxis)^2,54,55^, substrate-bound chemicals (haptotaxis)^2,56,57^, and the environmental stiffness (mechanosensing)^6,58^. Recent 2D analyses have shown that the adhesion-clutch system involving shootin1a mediates not only chemotaxis^25^, but also haptotaxis^26,59^ and mechanosensing^60^. During haptotaxis, differential mechanical interactions between adhesion molecules on growth cones and their adhesive ligands navigates axon pathfinding^26,59^. On the other hand, environmental stiffness modulates their interaction for mechanosensing^60^. We consider that such dynamic and specific adhesions between the growth cone and its environment enable 3D axon outgrowth, pathfinding and regeneration. In conclusion, the functionality of the adhesion-clutch machinery in a 3D environment is shown in this study, establishing it as a fundamental growth cone machinery in neural network formation.

## RESOURCE AVAILABILITY

### Lead Contact

Request for further information, resource, and reagents should be directed to and will be fulfilled by the lead contact, Naoyuki Inagaki.

### Materials Availability

All unique materials generated in this study are available from the lead contact with a completed Materials Transfer Agreement.

### Data and Code Availability

- All data reported in this paper will be shared by the lead contact upon request.
- Source data of immunostaining, immunoblot and statistical analyses in the figures are available at Mendeley Data (https://data.mendeley.com/datasets/rbs58jv2ps/3) and are publicly available as of the date of publication.
- Any additional information required to reanalyze the data reported in this paper is available from the lead contact upon request.

## ACKNOWLEDGMENTS

We thank Dr. Masatoshi Takeichi (Riken) for providing the N-cadherin cDNA; Mieko Ueda and Kazumi Maekawa for technical support; and Satoko Shimamura and Eisuke Inagaki for kind supports. This research was supported in part by AMED under Grant Number JP17gm0810011 (N.I.), JSPS KAKENHI (25K02272, N.I.), JSPS Grants-in-Aid for Early-Career Scientists (JP23K14181, T.M.), the Osaka Medical Research Foundation for Intractable Diseases (T.M.), and the NAIST Life Science Collaboration Center (LiSCo).

## AUTHOR CONTRIBUTION

H.C., Z.Q., S.Y, A.M., T.M., and N.I. designed the experiments. H.C., Z.Q., S.Y., and A.M., performed experiments and data analysis. N.I., H.C., Z.Q., and T.M. wrote the manuscript. N.I. supervised the project. All authors discussed the results and commented on the manuscript.

## SUPPLEMENTAL INFORMATION

Supplemental Information can be found online at XXX.

## STAR METHODS

### KEY RESOURCES TABLE

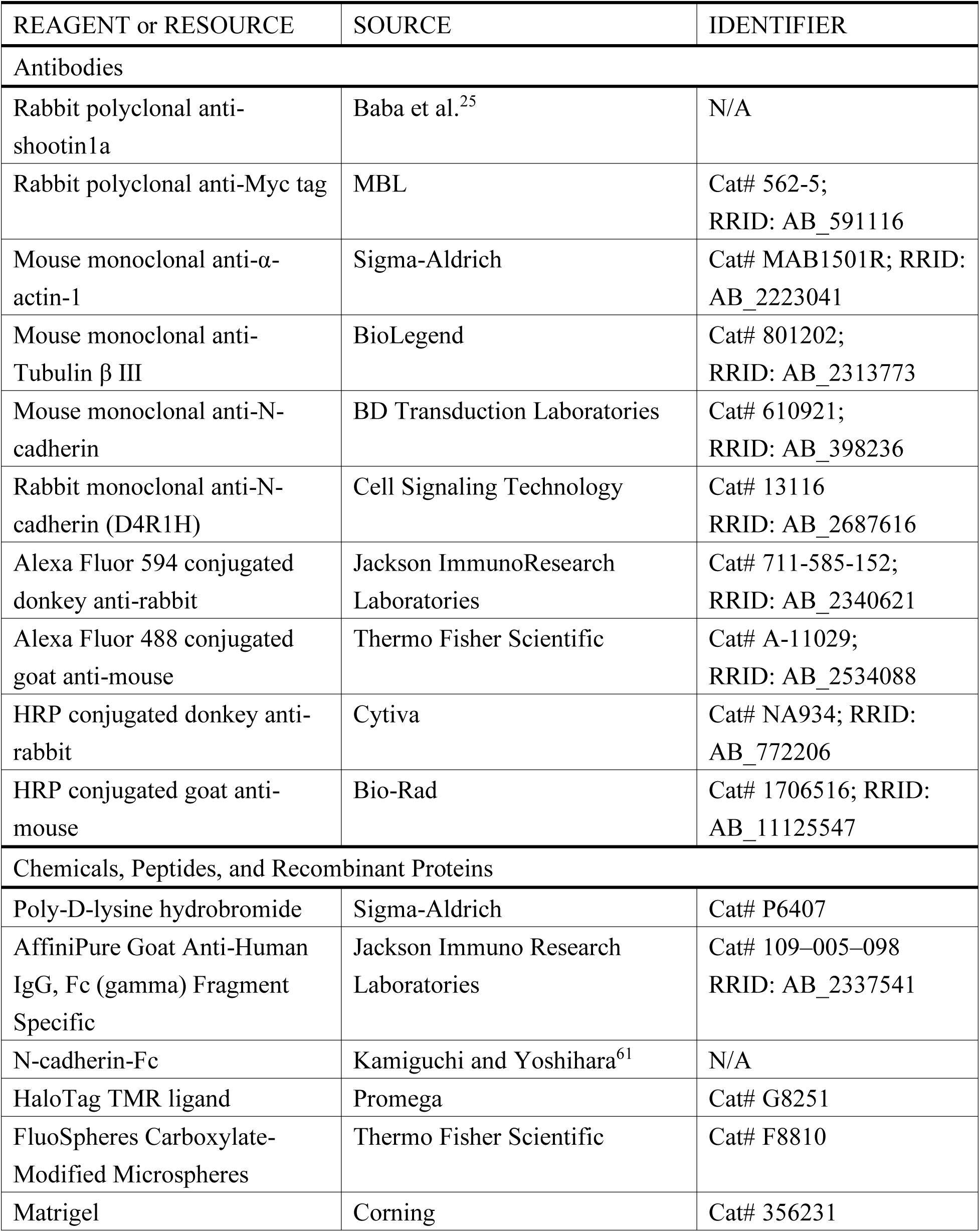
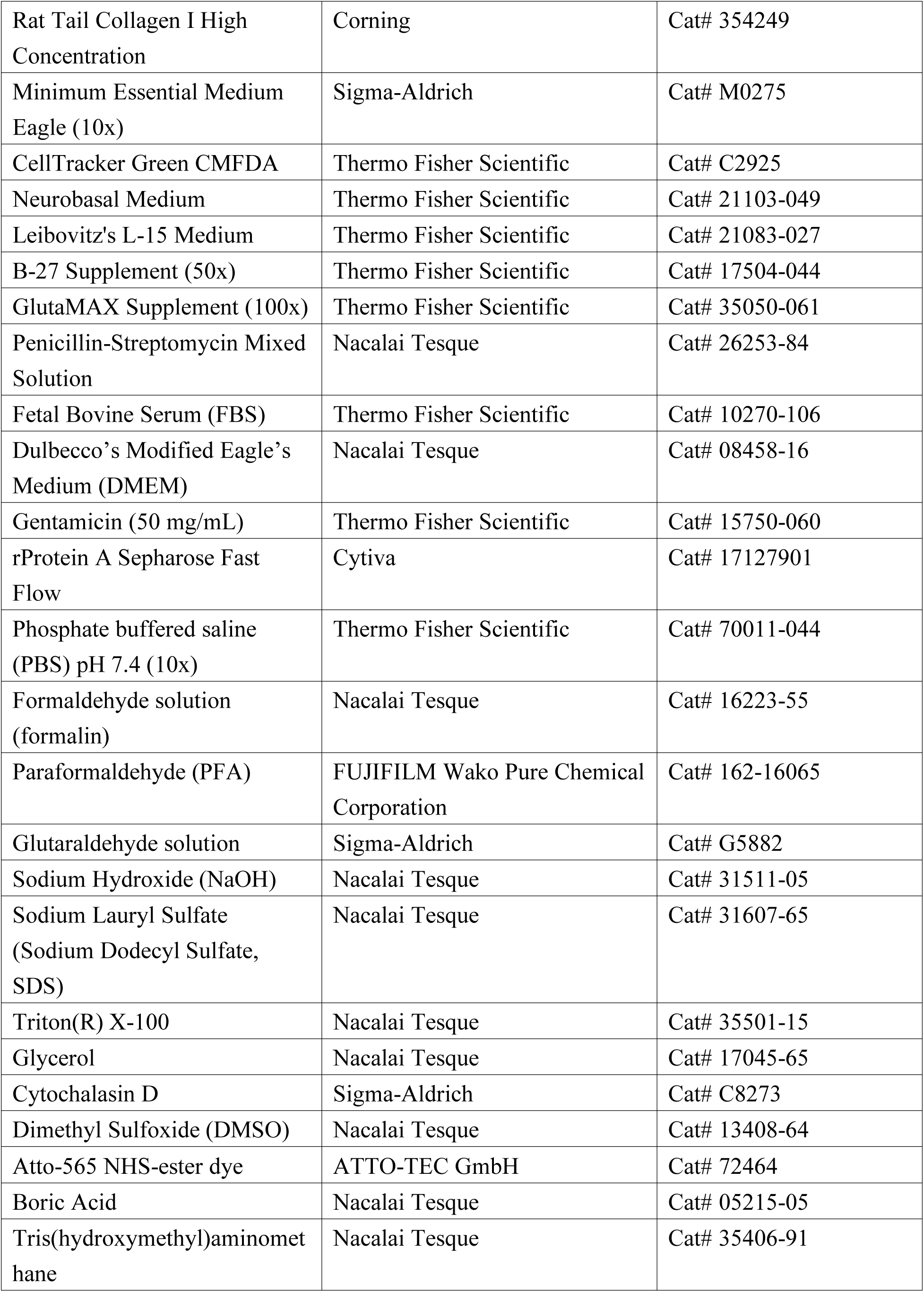
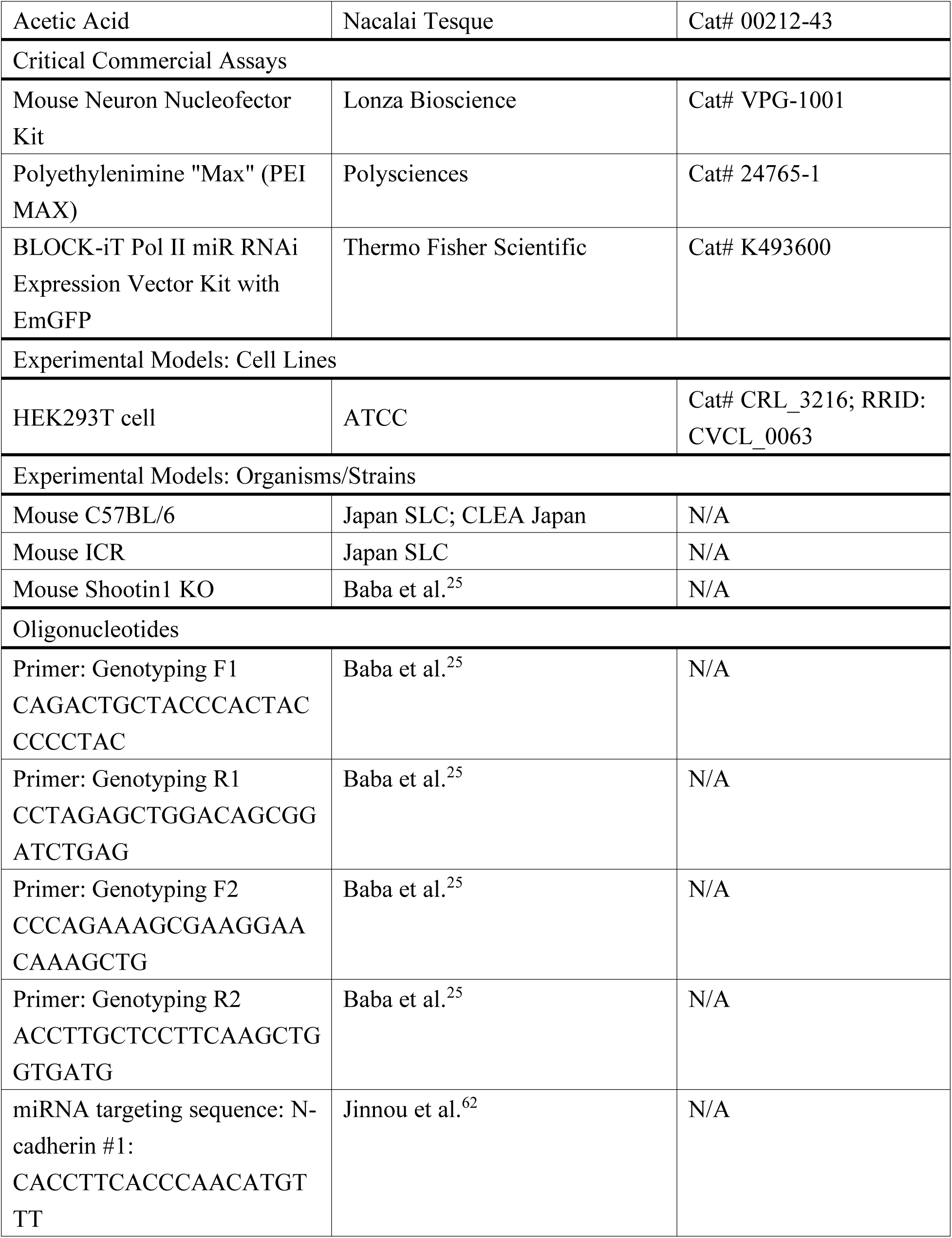
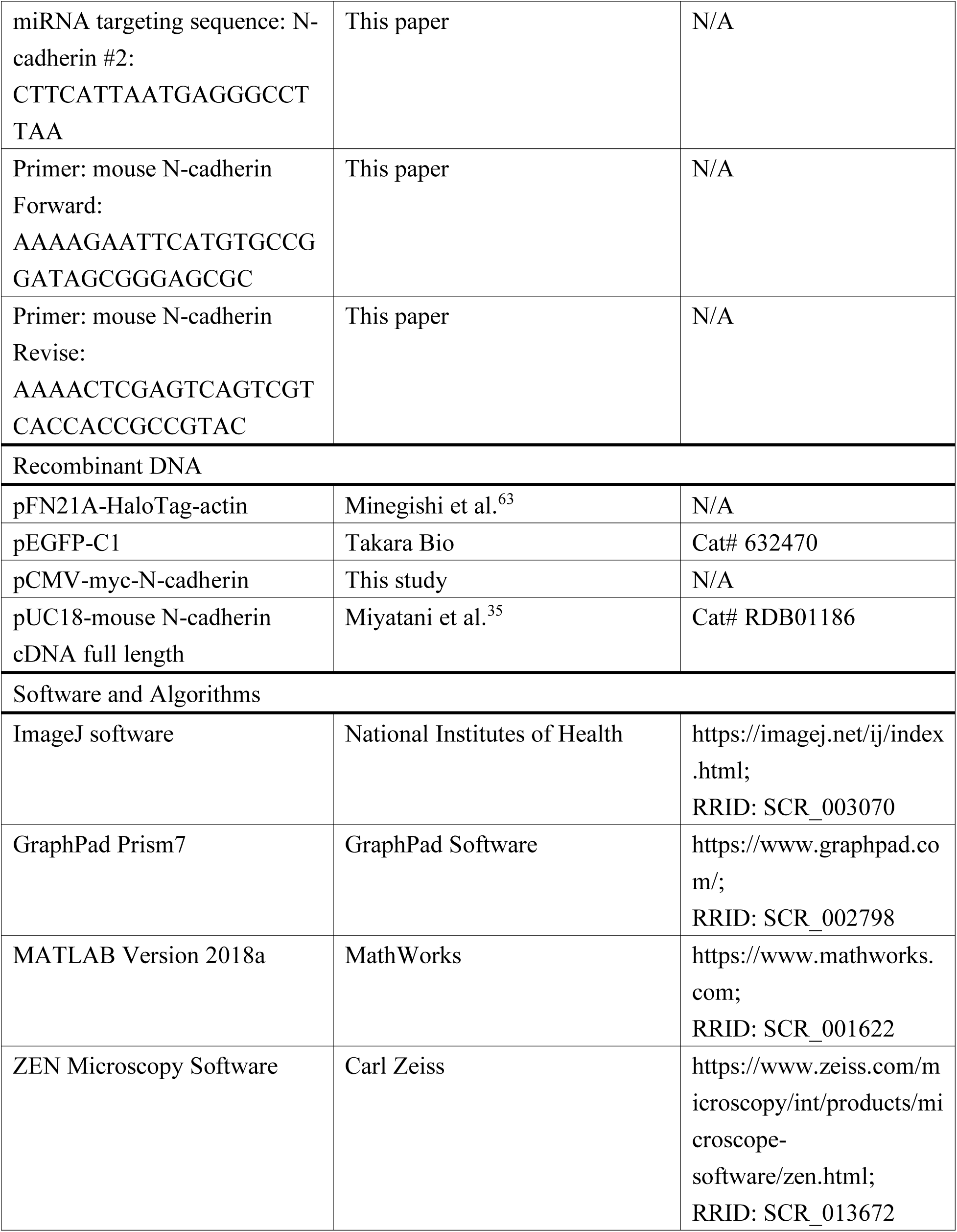
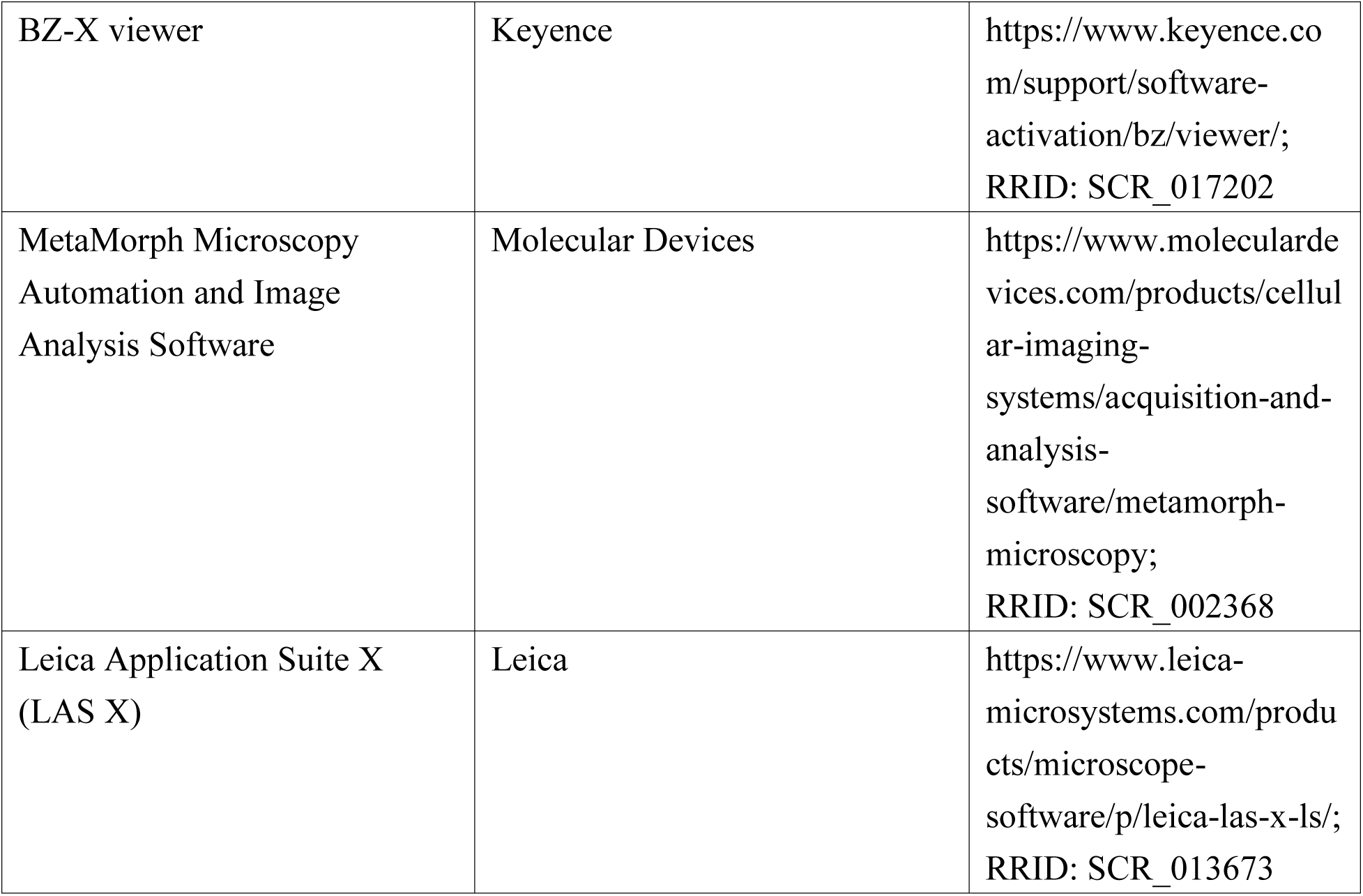

### EXPERIMENTAL MODEL AND SUBJECT DETAILS

#### Animals

All relevant aspects of the experimental procedures were approved by the Institutional Animal Care and Use Committee of Nara Institute of Science and Technology. Embryonic day 16.5 (E16.5) C57BL/6 and ICR mice were obtained from Japan SLC and CLEA Japan. Shootin1 KO mice were generated as described previously^25^. E16.5 shootin1 KO embryos carrying a homozygous mutation in the shootin1 gene were obtained by crossing male and female shootin1 heterozygous C57BL/6 mice; the offspring genotypes were checked by polymerase chain reaction (PCR) using the following primers: Genotyping F1 (5′-CAGACTGCTACCCACTACCCCCTAC-3′) and Genotyping R1 (5′-CCTAGAGCTGGACAGCGGATCTGAG-3′) for the wild-type (WT) allele. Genotyping F2 (5′-CCCAGAAAGCGAAGGAACAAAGCTG-3′), Genotyping R2 (5′-ACCTTGCTCCTTCAAGCTGGTGATG-3′) for the KO allele. Embryonic day 0.5 (E0.5) was defined as noon of the day when a vaginal plug was observed after overnight mating. Chimeric mice were crossed with C57BL/6 mice for at least nine generations before analysis. All C57BL/6 mice were bred under standard conditions (12 h/12 h light/dark cycle, access to dry food and water).

#### Transfection and cell culture in 2D condition

Hippocampal neurons prepared from E16.5 WT and shootin1 KO mouse embryos were cultured as described previously^29^.

For the experiments in Figures 1A, 1C, 1D, 2A-D, S1C, and S1E, neurons were seeded on 13-mm glass coverslips (Matsunami), 14-mm glass bottom dishes (Matsunami, D11130H) or polyacrylamide gels coated with 100 μg/mL poly-D-lysine (Sigma Aldrich, P6407) or coated sequentially with 100 μg/mL polylysine, 20 μg/µL anti-Fc antibody (Jackson Immuno Research Laboratories, 109-005-098) and 10 μg/mL N-cad-Fc^27,61^. Cells were subsequently cultured until DIV 2 in the culture medium [Neurobasal Medium (Thermo Fisher Scientific, 21103-049) containing 2% B-27 supplement (Thermo Fisher Scientific, 17504-044), 1% GlutaMAX supplement (Thermo Fisher Scientific, 35050-061) and 100 U/mL Penicillin-100 µg/mL streptomycin (Nacalai Tesque, 26253-84)] in a humidified incubator at 37 °C with 5% CO_2_.

For the experiments in Figures 2A-D, 1 × 10^6^ neurons were suspended in 100 μL of mouse neuron nucleofector solution (Lonza Bioscience, VPG-1001) and subsequently electroporated with vectors using an electroporation apparatus, Nucleofector 2b (Lonza Bioscience) before plating. Transfected cells were then seeded on polylysine or N-cad-Fc-coated glass bottom dishes or polyacrylamide gels. Then they were cultured until DIV 2 as described above.

HEK293T cells (ATCC, CRL-3216) were cultured in Dulbeco’s modified Eagle’s medium (DMEM) (Nacalai Tesque, 08458-16) supplemented with 10% fatal bovine serum (FBS) (Thermo Fisher Scientific, 10270-106), 100 U/mL Penicillin-100 µg/mL streptomycin and 0.01% gentamicin (Thermo Fisher Scientific, 15750-060) as described previously^25^. HEK293T cells were transfected with vectors using Polyethylenimine “Max” (PEI MAX) (Polysciences, 24765-1) following the manufacture’s protocol.

#### Transfection and cell culture in 3D Matrigel

For the experiments in Figures 3A, 4A, 4C, and S1F, neurons dissociated from E16.5 mouse hippocampi were suspended in phosphate buffered saline (PBS) (Thermo Fisher Scientific, 70011-044) in the presence or absence of N-cad-Fc (10 ng/µL). The cell suspension was then mixed with Matrigel (Corning, 356231) at a ratio of 1:3 and gently mixed (avoid bubbles) on ice. Twenty microliters of the mixture were immediately placed on the center of pre-cooled 14-mm glass bottom dishes and smeared evenly using a pipet tip. Then, the dishes were incubated in a humidified incubator at 37 °C with 5% CO_2_ for 3-5 min to allow Matrigel matrix to solidify, after which the gels were subsequently covered with the culture medium.

For the experiments in Figure 3A, cells were transfected with pFN21A-HaloTag-actin using Nucleofector^29^. The transfected cells were incubated with pre-warmed culture medium for 30 min and then mixed with Matrigel matrix as described above.

#### Cell culture in 3D collagen gel

For the experiments in Figure 4E and F, dissociated neurons were suspended in the culture medium and mixed with a collagen solution at a seeding density of 2.5 × 10^4^ cells per dish as previously described^32,64^ with modifications. The collagen solution was prepared by mixing rat tail collagen I (Corning, 354249, 10.4 mg/mL in 0.02N acetic acid; final concentration: 2.2 µg/µL) with PBS in the presence or absence of N-cad-Fc (final concentration: 2.5 ng/µL), 10x Minimum Essential Medium (MEM) (Sigma-Aldrich, M0275) (1:10 dilution) and 1 M NaOH (neutralizing solution; Nacalai Tesque, 31511-05) (final concentration: ∼0.0143 M). Cell suspension was then gently mixed with the collagen solution at a ratio of 1:4 (avoid bubbles). Fifty microliters of the mixture were immediately transferred to 14-mm glass bottom dishes and smeared evenly with pipet tips. Then, the dishes were incubated in a humidified incubator at 37 °C with 5% CO_2_ for 30 min for polymerization of collagen fibers, after which the gels were subsequently covered with the culture medium.

#### Cell culture in 3D Matrigel containing fluorescently labeled collagen fibers

For the experiment in Figure 3D, neurons were suspended in the culture medium and mixed with Matrigel containing fluorescently labeled collagen fibers and N-cad-Fc at a seeding density of 2.5 × 10^4^ cells per dish as previously described^32,33^ with modifications. The collagen solution was prepared by mixing fluorescently labelled collagen (2.5 mg/mL) with N-cad-Fc diluted in PBS (final concentration: 2.5 ng/µL), 10x MEM (1:10 dilution) and 1 M NaOH; final concentration: ∼0.0143 M). After adjusting the pH to ∼7-7.4 with NaOH, the fluorescently labelled collagen solution was then gently mixed with Matrigel matrix at a ratio of 5:3 (containing ∼ 37.5% Matrigel) on ice^32^. The cell suspension was immediately mixed with the Matrigel-collagen solution at a proportion of 1:3 and gently mixed on ice. Forty microliters of the mixture were plated on the center of each 14-mm glass bottom dish and smeared evenly by a pipet tip. Then dishes were incubated at 37 °C with 5% CO_2_ for 30 min for gelation of 3D hydrogel, after which the gels were subsequently covered with the culture medium.

### METHOD DETAILS

#### Drug treatments

Cytochalasin D (Sigma-Aldrich, C8273), dissolved in DMSO (Nacalai Tesque, 13408-64), was added to the culture medium at a final concentration of 1 µM. The same volume of DMSO at a dilution of 1:1,000 was added to the medium as a control. For the experiment in Figure S1C, neurons were treated with 1 μM cytochalasin D or 0.1% DMSO after plating, and then cultured for 40 h until DIV 2. For the experiments in Figures 4C, 4E and 4F, cytochalasin D treatment was carried out when the culture medium was added on 3D Matrigel or collagen gel.

#### DNA construction and RNAi

Preparation of pFN21A-HaloTag-actin was described previously^63^.

The full-length cDNA of mouse N-cadherin was provided by the RIKEN BRC through the National BioResource Project of the MEXT, Japan (cat. RDB01186)^35^. To generate pCMV-myc-N-cadherin, mouse N-cadherin cDNA was amplified by PCR with the primers 5′-AAAAGAATTCATGTGCCGGATAGCGGGAGCGC-3’ and 5’-AAAACTCGAGTCAGTCGTCACCACCGCCGTAC-3’. Mouse N-cadherin cDNA was then subcloned into pCMV-myc vector (fused to N-terminal myc-tag, Agilent Technologies) with a cytomegalovirus (CMV) promoter.

For vector-based mouse N-cadherin RNAi experiment, we used a BLOCK-iT Pol II miR RNAi Expression Vector Kit with EmGFP (Thermo Fisher Scientific, K493600) and Block-iT RNAi designer. The targeting sequences of N-cadherin miRNA #1 (5′-CACCTTCACCCAACATGTTT-3′) was reported previously^62^, which corresponds to nucleotides 944-963 in the coding region of mouse N-cadherin and was inserted into the pcDNA6.2-GW/EmGFP-miR expression vector. The targeting sequences of N-cadherin miRNA #2 (5′-CTTCATTAATGAGGGCCTTAA-3′) corresponds to nucleotides 3354–3374 in the coding region of mouse N-cadherin. The control vector pcDNA 6.2-GW/EmGFP-miR-neg encodes a miRNA (5′-GAAATGTACTGCGCGTGGAGACGTTTTGGCCACTGACTGACGTCTCCACGCAGTACATTT-3′) that targets no known vertebrate gene. These miRNA expression vectors are designed to co-express EGFP.

As described previously^17^, to ensure a high-level expression of N-cadherin miRNA before axon outgrowth, hippocampal neurons transfected with the miRNA expression vector [control miRNA or N-cadherin miRNA (#1)], using Nucleofector 2b, were cultured in suspension onto uncoated polystyrene dishes. After a 24-h incubation to induce miRNA expression, cells were suspended in the culture medium (Figure S1C) or PBS containing 10 ng/µL N-cad-Fc (Figure S1F), and then cultured on N-cad-Fc-coated coverslips (Figure S1C) or in 3D Matrigel (Figure S1F) for 40 h until DIV 2 as described above.

To confirm the reduction of mouse N-cadherin expression levels (Figurs S1B), HEK293T cells were co-transfected with pCMV-myc-N-cadherin and miRNA expression vector [control miRNA or N-cadherin miRNA (#1 or #2)] using PEI MAX. After 48 h, transfected cells were collected and lysed with RIPA buffer [50 mM Tris-HCl (pH 8.0), 1 mM EDTA, 150 mM NaCl, 1% Triton X-100 (Nacalai Tesque, 35501-15), 0.1% sodium dodecyl sulfate (SDS, Nacalai Tesque, 31607-65), 0.1% sodium deoxycholate, 1 mM DTT, 1 mM PMSF, and 0.01 mM leupeptin] on ice for 10 min. The cell lysate was then centrifuged at 17,900 × g for 10 min at 4 °C. The supernatant was mixed with an equal volume of 2 x SDS sample buffer (131 mM Tris-HCl, pH 6.8, 21% glycerol, 4% SDS, 12 M urea, 0.05% bromophenol blue and 5% β-mercaptoethanol) and incubated for 5 min at 95 °C. Immunoblot was performed as described previously^25^. The following primary antibodies were used in immunoblot: rabbit anti-Myc tag (1:2000) (MBL, 562-5) and mouse anti-actin (1:5,000) (Sigma-Aldrich, MAB1501R) antibodies. The following secondary antibodies were used for immunoblot: HRP-conjugated donkey anti-rabbit IgG (1:2,000) (Cytiva, NA934) and HRP-conjugated goat anti-mouse IgG (1:4,000) (Bio-Rad, 1706516) antibodies.

#### Preparation of purified N-cadherin-Fc

N-cadherin-Fc was prepared as described previously^61^. Briefly, HEK293T cells were transfected with the vector expressing N-cadherin-Fc^61^ using PEI MAX. After 6 h, the culture medium was replaced with serum-free DMEM containing 100 U/mL Penicillin-100 µg/mL streptomycin and 0.01% gentamicin. The serum-free medium was conditioned for 3-4 days. N-cad-Fc was purified from the culture supernatant using a rProtein A Sepharose Fast Flow column (Cytiva, 17127901) and dialyzed with PBS containing 0.9 mM CaCl_2_ and 0.9 mM MgCl_2_.

#### Fluorescent labeling of collagen

Fluorescently labelled collagen solution was prepared as described^65^. Briefly, 5 mL of rat tail collagen I was polymerized at room temperature for 30 min. After polymerization, the collagen gel was incubated with 50 mM boric acid (pH 9.0) (Nacalai Tesque, 05215-05) for 10 min. Then the borate buffer was removed, and the gel was incubated with 5 mL of Atto-565 NHS-ester dye solution (ATTO-TEC GmbH, 72464) in borate buffer at 4°C overnight in the dark. The concentration of dye (diluted in DMSO) was adjusted to a 2-molar excess as recommended by AttoTech. The dye solution was subsequently removed, and the remaining Atto-565 NHS-ester dye was quenched with 10 mL of 50 mM Tris buffer (pH 7.4) (Nacalai Tesque, 35406-91) for 10 min. The gels were then rinsed 6-10 times with PBS for 6-10 h. Then the gel was acidified in 1 mL of 500 mM acetic acid (Nacalai Tesque, 00212-43) and stirred till the gel was completely solubilized. The solution was then dialyzed in 20 mM acetic acid at 4°C for 24 h. The final concentration of collagen was determined by SDS-PAGE and adjusted at 2.5 mg/mL.

#### Immunocytochemistry and microscopy

Neurons cultured on 2D glass substrates were fixed with 3.7% formaldehyde (Nacalai Tesque, 16223-55) diluted in Krebs buffer (118 mM NaCl, 4.7 mM KCl, 1.2 mM KH_2_PO_4_, 1.2 mM MgSO_4_, 4.2 mM NaHCO_3_, 2 mM CaCl_2_, 10 mM glucose, 400 mM sucrose, 10 mM HEPES; pH 7.0) for 5 min at room temperature and then placed on ice for 20 min. Neurons cultured in 3D collagen gel or Matrigel were fixed with 4% (w/v) paraformaldehyde (PFA) (FUJIFILM Wako Pure Chemical Corporation, 162-16065) and 1% (w/v) glutaraldehyde (Sigma-Aldrich, G5882) in PBS for 30 min at 37°C.

For the experiments in Figures 1A, 1C, 1D and S1E, the fixed cells were permeabilized with 0.05% Triton X-100 in PBS for 15 min on ice and 10% FBS in PBS for 1 h at room temperature. Neurons were then incubated with primary antibody diluted in PBS (containing 10% FBS) overnight at 4°C. The following primary antibodies were used: rabbit anti-shootin1a^25^ (1:1,000), mouse monoclonal anti-N-cadherin-ICD (1:1,000) (BD Transduction Laboratories, 610921), and mouse monoclonal anti-Tubulin β Ⅲ (TUJ1) (1:1,000) (BioLegend, 801202) antibodies. After washing with PBS, cells were immunostained with secondary antibody diluted in PBS at RT for 1 h. The following secondary antibodies were used: Alexa Fluor 594 conjugated donkey anti-rabbit (1:1,000) (Jackson ImmunoResearch Laboratories, 711-585-152) and Alexa Fluor 488 conjugated goat anti-mouse (1:1,000) (Thermo Fisher Scientific, A-11029) antibodies. After washing with PBS, immunostained neurons were mounted with 50% (v/v) glycerol (Nacalai Tesque, 17045-65) in PBS.

For the experiments in Figures 4E and 4F, the fixed cells were treated with 0.4% Triton X-100 in PBS for 30 min at room temperature and then incubated with the blocking solution (PBS containing 10% FBS and 0.4% Triton X-100) for 1 h at room temperature. Cells were then immunostained with mouse monoclonal anti-Tubulin β Ⅲ antibody (1:1,000) diluted in the blocking solution overnight at 4°C. After washing with 0.4% Triton X-100 in PBS, cells were incubated with Alexa Fluor 488 conjugated goat anti-mouse antibody (1:1,000) diluted in PBS (containing 0.4% Triton X-100) at RT for 1 h.

For the microscopy in Figures 1A, 1D, 4A, 4C, S1C, S1E and S1F, fluorescence or brightfield images were acquired using an inverted fluorescence phase contrast microscope (BZ-X710, Keyence) equipped with a Plan APO VC 20×/0.75 NA Air objective lens (Nikon, 0500-0087) and imaging software (BZ-X viewer, Keyence). To acquire images of the entire hippocampal neurons for the microscopy in Figure S1F, we obtained ∼10-50 images at 0.5 μm intervals along the Z-axis, which were then volume-stacked. Axon length was measured using ImageJ (National Institutes of Health).

For the microscopy in Figure 1C, fluorescence images were captured by a total internal reflection fluorescence (TIRF) microscope (IX81, Olympus) equipped with an EM-CCD camera (Ixon3, Andor), a UAPO N 100×/1.49 NA Oil Immersion OTIRF Apochromat objective lens (Olympus) and imaging software (MetaMorph Microscopy Automation and Image Analysis Software, Molecular Devices).

For the microscopy in Figure 4E and F, fluorescence images were obtained using a confocal laser microscope (Stellaris 8, Leica) equipped with a HC PL APO 40x/1.30 OIL CS2 objective lens (Leica, 11506428) and imaging software (Leica Application Suite X, Leica). To acquire images of the entire hippocampal neurons, we obtained ∼40-170 confocal images at 0.5 μm intervals along the Z-axis. The confocal images were then volume-stacked and the axon length was quantified using ImageJ.

#### Immunostaining of 3D hydrogel

For the experiments in Figures 3B and 4D, the hydrogels were fixed with 4% (w/v) PFA and 1% glutaraldehyde in PBS for 30 min at 37°C. The gels were then treated with the blocking solution (PBS containing 10% FBS and 0.4% Triton X-100) for 1 h at room temperature. The gels were then incubated with mouse monoclonal anti-N-cadherin antibody (D4R1H, Cell Signaling Technology, 13116) (1:200) diluted in the blocking solution overnight at 4°C. After washing with 0.4% Triton X-100 in PBS, they were incubated with Alexa Fluor 488 conjugated goat anti-mouse antibody (1:1,000) diluted in PBS (containing 0.4% Triton X-100) at RT for 1 h. After washing with 0.4% Triton X-100 in PBS, gels were further washed with PBS and observed using a confocal laser microscope (LSM980, Carl Zeiss) equipped with a C-Apochromat 63x/1.2 W Corr objective lens (Carl Zeiss) and imaging software (ZEN Microscopy Software, Carl Zeiss).

#### Fluorescent speckle imaging

Fluorescent speckle imaging and speckle tracking analysis of HaloTag-actin was performed as described previously^29^. The plasmid to express HaloTag-actin was transfected into hippocampal neurons using Nucleofector 2b. The fluorescently labeled actin molecules replace some of the endogenous actins to form F-actins and move retrogradely in axonal growth cones, depicting the F-actin movement. Prior to observation, neurons were treated with HaloTag TMR ligand (Promega, G8251) diluted in the culture medium (at a final concentration of 50 nM) and incubated for 1 h at 37°C with 5% CO_2_. The ligand was then washed out with pre-warmed PBS, and cells were incubated with the pre-warmed observation medium [Leibovitz’s L-15 Medium (Thermo Fisher Scientific, 21083-027) containing 2% B27 supplement, 1% GlutaMAX supplement and 100 U/mL Penicillin-100 µg/mL streptomycin]. The fluorescent speckles of HaloTag-actin were observed using a fluorescence microscope (AxioObserver Z1, Carl Zeiss) equipped with a complementary metal oxide semiconductor (CMOS) camera (ORCA Flash4.0 V2, Hamamatsu), a Plan-Apochromat 100x/1.4 Oil DIC M27 objective lens (Carl Zeiss), and imaging software (ZEN Microscopy Software, Carl Zeiss). Fluorescence images were acquired every 5 s. The speed of F-actin flow was determined by tracking fluorescent actin speckles using ImageJ. Briefly, filopodium consist of clear fluorescent actin speckles at the leading edge of axonal growth cone was selected and a time-lapse montage was made to show the position of each speckle at different time-point. Clear speckles moved retrogradely for at least 15 s (3 frames) were used for analysis. The velocity of actin retrograde movement was calculated through dividing the speckle translocation distance by the observation time.

#### Traction force microscopy

Traction force microscopy in Figures 2C and 2D was performed as previously described^29^. Hippocampal neurons were electroporated with pEGFP-C1 vector (Takara Bio, 632470) using Nucleofector 2b and then cultured until DIV2 on polylysine or N-cad-Fc-coated polyacrylamide gels with embedded 200-nm fluorescent microspheres (Thermo Fisher Scientific, F8810). Prior to observation, culture medium was replaced with the pre-warmed observation medium. Time-lapse fluorescence and differential interference contrast (DIC) images were acquired every 3 s at 37°C using a confocal laser microscope (LSM710, Carl Zeiss) equipped with a C-Apochromat 63x/1.2 W Corr objective lens (Carl Zeiss) and imaging software (ZEN Microscopy Software, Carl Zeiss). The growth cone areas of transfected neurons were identified by both EGFP fluorescence and DIC images. After imaging, culture dishes were treated with 10% (w/v) SDS to release neurons from the polyacrylamide gel. The image of the fluorescent beads in the unstrained substrate was captured and used as a reference for the original bead positions. The magnitude of traction force under the axonal growth cone was determined by monitoring the fluorescent bead displacement from their original position using MATLAB (MathWorks) and a homemade algorithm reported previously^29^.

For the experiments in Figure 3D, neurons were cultured in 3D Matrigel containing fluorescently labeled collagen fibers and N-cad-Fc until DIV2. Prior to observation, neurons were stained with a volume marker, CMFDA (Thermo Fisher Scientific, C2925) diluted in the culture medium (at a final concentration of 200 nM) and incubated for 1 h at 37°C. The cells were then washed with pre-warmed PBS and incubated with the pre-warmed observation medium. CMFDA-labeled axonal growth cones and 3D hydrogel were observed using a confocal laser microscope (Stellaris 8, Leica) equipped with a HC PL APO 100x/1.40 OIL CS2 objective lens (Leica, 11506372) and imaging software (LAS X, Leica). Time-lapse fluorescence images were obtained every 10 s with a Z-stack of 3-5 μm at 0.5 μm intervals and the time-lapse images were volume-stacked. After imaging, the culture dishes were treated with 2% Triton X-100 to lyse the neurons from the gel. The fluorescence image of the unstrained gel was acquired with a Z-stack of 3-5 μm at 0.5 μm intervals. The images were then volume-stacked and used as a reference for the original fiber positions.

### QUANTIFICATION AND STATISTICAL ANALYSIS

All statistical analyses were performed using Excel 2016 (Microsoft) and GraphPad Prism7 (GraphPad Software). The D’Agostino–Pearson normality test was used to determine whether the data followed a normal distribution. We also tested the equality of variation with the F-test for two independent groups that followed normal distributions. Statistical significance tests were performed as following: (1) two-tailed paired *t*-test for the comparison between two dependent groups that showed normal distribution; (2) two-tailed unpaired Student’s *t*-test for the comparison between two independent groups that showed normal distribution and equal variation; (3) two-tailed unpaired Welch’s *t*-test for the comparison between two independent groups that showed normal distribution and unequal variation; (4) Mann–Whitney *U*-test for the comparison between two independent groups that showed non-normal distribution. For each experiment, the corresponding statistical information and number of samples are indicated in figure legends. All data are shown as the mean ± SEM. Statistical significance was defined as ***p < 0.01; **p < 0.02; *p < 0.05; ns, not significant. P value less than 0.05 was considered to be statistically significant. All experiments were performed at least three times and reliably reproduced. Investigators were blind to experimental groups for each analysis, except biochemical analysis.

To quantify the degree of colocalization between shootin1a and N-cadherin, the immunostaining data in Figure S1D were analyzed by Pearson’s correlation coefficient (PCC) using JaCoP^66^, a plugin of ImageJ. PPC values between 0.5 and 1 represent the colocalization of two fluorescent signals.

