## Supplemental Figure 1 for "Adhesion-clutch drives three-dimensional axon outgrowth"

**This PDF file includes:**

SUPPLEMENTAL DISCUSSION

Figure S1

Supplemental Video legends

**Other supplemental information for this manuscript include:**

Supplemental Videos S1-S6

**SUPPLEMENTAL DISCUSSION**

This section provides additional discussion of the results that were not covered in the main text.

1) Figures 1A and 1B show that the axon length of neurons cultured on N-cad-Fc in Figure 1B (control RNAi) is shorter than that in Figure 1A (N-cad-FC (+)). In Figure 1A, dissociated hippocampal neurons were cultured on N-cad-Fc-coated coverslips (N-cad-Fc (+)) for 40h. In Figure 1B, neurons were transfected with the control miRNA expression vector (control miRNA) and cultured in suspension in uncoated polystyrene dishes. After a 24-h incubation to induce miRNA expression, the cells were seeded on N-cad-Fc-coated coverslips and cultured for 40 h. We think that the difference is derived from the different culture conditions in Figures 1A and 1B.
2) Figures 1D and 1E show that the axon length of WT neurons in Figure 1E (control) is shorter than that in Figure 1D (WT). The culture medium of the experiment in Figure 1E contains 0.1% DMSO (vehicle) that inhibits neurite outgrowth[1]. We think that DMSO in the experiment in Figure 1E inhibited axon outgrowth of both control and cytochalasin D-treated neurons.
3) Figure 4A shows that depletion of shootin1a led to stronger inhibition of axon outgrowth than N-cad-Fc depletion, and similar effects were observed in F-actin retrograde flow in Figure 3A. We previously reported that the adhesion-clutch system involving shootin1a and the cell adhesion molecule L1 drives 2D axon outgrowth[2]. As Matrigel contains the L1 ligand laminins[2], We consider that shootin1a KO inhibits both N-cadherin-shootin1a adhesion-clutch and L1-shootin1a adhesion-clutch in 3D Matrigel. On the other hand, the N-cad-Fc depletion inhibits only the N-cadherin-shootin1a adhesion-clutch. Therefore, the data in Figures 4A and 3A suggest that L1-shootin1a adhesion-clutch also works in 3D environment in the presence of its adhesive ligand laminin.

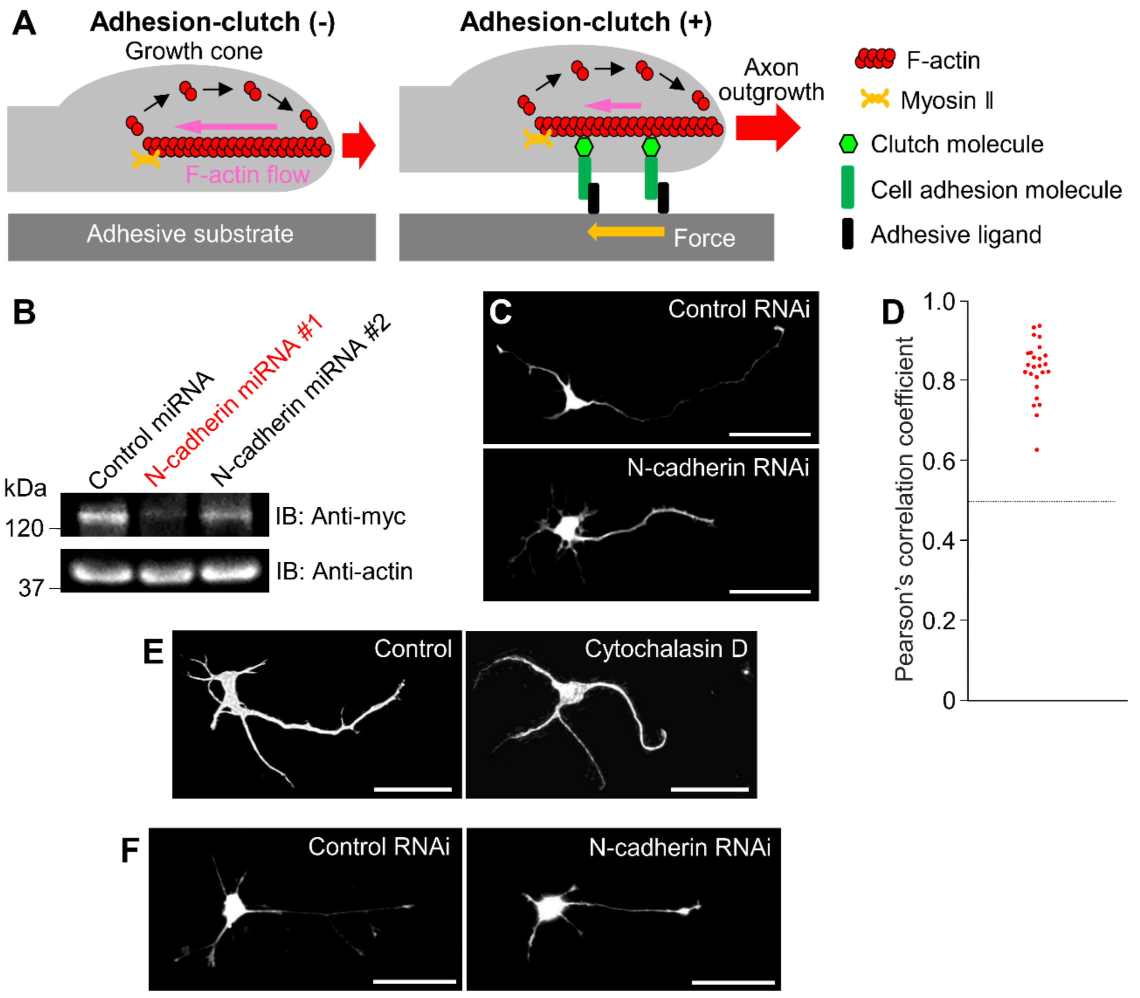


**Figure S1. N-cadherin and shootin1a form adhesion-clutch for 2D and 3D axon outgrowth, related to Figure 1 and 4**

(A) A mechanical model (adhesion-clutch mechanism) explaining the traction force generated by axonal growth cones during forward migration in 2D environments. Actin filaments (F-actins) undergo continuous retrograde flow in axonal growth cones, which is powered by the combination of actin polymerization at the leading edge, the proximal depolymerization, and the contractile activity of myosin II. Neuronal clutch molecules (such as shootin1a) and cell adhesion molecules (such as N-cadherin) cooperatively mediate the mechanical coupling between the F-actin flow and extracellular adhesive substrates. Such coupling reduces the speed of actin retrograde flow (pink arrows) and transmits the rearward movement of treadmilling actin filaments into the backward traction force (yellow arrows), whose counter force (pulling force, red arrows) is required for 2D axon outgrowth. Loss of clutch molecules or cell adhesion molecules increases the speed of F-actin flows (pink arrows) and decreases the traction force generation (yellow arrows), therefore inhibiting the axon outgrowth in 2D environments.

B) HEK293T cells were co-transfected with mouse pCMV-myc-N-cadherin vector and candidate mouse N-cadherin RNAi vectors (#1 or #2) or control miRNA vector. Cell lysates were analyzed by immunoblot with anti-myc tag antibody. Actin was used as a loading control. N-cadherin miRNA vector #1 was used to knockdown endogenous N-cadherin in hippocampal neurons as shown in Figures 1B, 4B, S1C and S2.

(C) Fluorescence images of hippocampal neurons transfected with the miRNA expression vector [(control miRNA or N-cadherin miRNA (#1), designed to co-express EGFP] and cultured in suspension onto uncoated polystyrene dishes for 24 h. Cells were then seeded on N-cad-Fc-coated coverslips for 40 h until DIV 2. Scale bar, 50 μm.

(D) Quantification of shootin1a-N-cadherin colocalization in DIV 2 axonal growth cone by Pearson’s coefficient correlation, a statistic method used to quantify the degree of colocalization between two fluorescent signals using JACoP (ImageJ plugin). The Pearson’s coefficient correlations of all growth cones exceeded 0.5 ($n = 25$ growth cones), indicating the colocalization of shootin1a and N-cadherin in axonal growth cones.

(E) Fluorescence images of hippocampal neurons cultured on N-cad-Fc-coated coverslips and treated with 0.1% DMSO (control) or 1 μM cytochalasin D for 40 h until DIV 2. Neurons were immunostained with anti-Tuj1 antibody. Scale bar, 50 μm.

(F) Fluorescence images of hippocampal neurons transfected with the miRNA expression [(control miRNA or N-cadherin miRNA (#1), designed to co-express EGFP] and cultured in suspension onto uncoated polystyrene dishes for 24 h. Cells were then embedded in 3D Matrigel containing N-cad-Fc for 40 h until DIV 2. Scale bar, 50 μm.

### SUPPLEMENTAL VIDEO LEGENDS

**Video S1. Fluorescent Speckle Imaging of actin dynamics in axonal growth cones of hippocampal neurons cultured on polylysine or N-cad-Fc, related to Figure 2A.**
Time-lapse fluorescence movies of HaloTag-actin in axonal growth cones of DIV 2 hippocampal neurons cultured on polylysine (N-cad-Fc (-)) or N-cad-Fc-coated (N-cad-Fc (+)) glass-bottom dishes. Images were acquired every 5 s for 295 s using a fluorescence microscope (AxioObserver Z1, Carl Zeiss).

**Video S2. Fluorescent Speckle Imaging of actin dynamics in axonal growth cones of WT and shootin1 KO hippocampal neurons cultured on N-cad-Fc, related to Figure 2B.**
Time-lapse fluorescence movies of HaloTag-actin in axonal growth cones of WT and shootin1 KO hippocampal neurons (DIV 2) cultured on glass-bottom dishes coated sequentially with polylysine, anti-Fc and N-cad-Fc. Images were captured every 5 s for 295 s using a fluorescence microscope (AxioObserver Z1, Carl Zeiss).

**Video S3. Traction Force Microscopy of axonal growth cones of hippocampal neurons cultured on polylysine or N-cad-Fc, related to Figure 2C.**
Time-lapse fluorescence movies of axonal growth cones of DIV 2 hippocampal neurons expressing EGFP (blue) and cultured on polylysine (N-cad-Fc (-)) or N-cad-Fc-coated (N-cad-Fc (+)) polyacrylamide gel embedded with 200-nm fluorescent beads. The original and displaced beads positions in the gel are indicated by green and red colors, respectively. White lines indicate the boundaries of axonal growth cones. Images of fluorescent beads and axonal growth cones were obtained every 3 s for 147 s using a confocal laser microscope (LSM710, Carl Zeiss).

**Video S4. Traction Force Microscopy of axonal growth cones of WT and Shootin1 KO hippocampal neurons cultured on N-cad-Fc, related to Figure 2D.**
Time-lapse fluorescence movies of axonal growth cones of WT and shootin1 KO hippocampal neurons (DIV 2) expressing EGFP (blue) and cultured on polyacrylamide gel embedded with 200-nm fluorescent beads and coated sequentially with polylysine, anti-Fc as well as N-cad-Fc. The original and displaced positions of the beads in the gel are indicated by green and red colors, respectively. White lines indicate the growth cone boundaries. Images of fluorescent beads and axonal growth cones were obtained every 3 s for 147 s using a confocal laser microscope (LSM710, Carl Zeiss).

**Video S5. Fluorescent Speckle Imaging of actin dynamics in axonal growth cones of WT and shootin1 KO hippocampal neurons cultured in 3D Matrigel, related to Figure 3A.**
Time-lapse fluorescence movies of HaloTag-actin in axonal growth cones of WT hippocampal neurons (DIV 2) cultured in 3D Matrigel added with (N-cad-Fc (-) WT) or without N-cad-Fc (N-cad-Fc (+) WT), and shootin1 KO neurons (DIV 2) cultured in Matrigel containing N-cad-Fc (N-cad-Fc (+) Shootin1 KO). Images were obtained every 5 s for 295 s using a fluorescence microscope (AxioObserver Z1, Carl Zeiss).

**Video S6. Traction Force Microscopy of an axonal growth cone of a hippocampal neuron cultured in 3D hydrogel containing N-cad-Fc, related to Figure 3D.**
Time-lapse fluorescence movies of an axonal growth cones of a DIV 2 hippocampal neurons cultured in 3D Matrigel containing fluorescently labeled collagen fibers and N-cad-Fc. The growth cone was labeled by the volume marker, CMFDA (blue). The original and displaced positions of collagen fibers are indicated by green and red colors, respectively. White line indicates the growth cone boundary. White dashed rectangle 1 indicated a fiber directly interacts with a filopodia. White dashed rectangle 2 indicated a fiber directly interacts with filopodium. Images were acquired every 10 s for 190 s at 0.5 µm intervals in the Z axis direction using a confocal laser microscope (Stellaris 8, Leica).
